# The genomic landscape at a late stage of stickleback speciation: high genomic divergence interspersed by small localized regions of introgression

**DOI:** 10.1101/190629

**Authors:** Mark Ravinet, Kohta Yoshida, Shuji Shigenobu, Atsushi Toyoda, Asao Fujiyama, Jun Kitano

## Abstract

Speciation is a continuous process and analysis of species pairs at different stages of divergence provides insight into how it unfolds. Genomic studies on young species pairs have often revealed peaks of divergence and heterogeneous genomic differentiation. Yet it remains unclear how localised peaks of differentiation progress to genome-wide divergence during the later stages of speciation with gene flow. Spanning the speciation continuum, stickleback species pairs are ideal for investigating how genomic divergence builds up during speciation. However, attention has largely focused on young postglacial species pairs, with little known of the genomic signatures of divergence and introgression in older systems. The Japanese stickleback species pair, composed of the Pacific Ocean three-spined stickleback (*Gasterosteus aculeatus*) and the Japan Sea stickleback (*G. nipponicus*), which co-occur in the Japanese islands, is at a late stage of speciation. Divergence likely started well before the end of the last glacial period and crosses between Japan Sea females and Pacific Ocean males result in hybrid male sterility. Here we use coalescent analyses and Approximate Bayesian computation to show that the two species split approximately 0.68-1 million years ago but that they have continued to hybridise at a low rate throughout divergence. Population genomic data revealed that high levels of genomic differentiation are maintained across the majority of the genome when gene flow occurs. However despite this, we identified multiple, small regions of introgression, strongly correlated with recombination rate. Our results demonstrate that a high level of genome-wide divergence can establish in the face of persistent introgression and that gene flow can be localized to small genomic regions at the later stages of speciation with gene flow.

**Author summary:** When species evolve, reproductive isolation leads to a build-up of differentiation in the genome where genes involved in the process occur. Much of our understanding of this comes from early stage speciation, with relatively few examples from more divergent species pairs that still exchange genes. To address this, we focused on Pacific Ocean and Japan Sea sticklebacks, which co-occur in the Japanese islands. We established that they are the oldest and most divergent known stickleback species pair, that they evolved in the face of gene flow and that this gene flow is still on going. We found introgression is confined to small, localised genomic regions where recombination rate is high. Our results show high divergence can be maintained between species, despite extensive gene flow.

## Introduction

Spéciation is a continuous process through which reproductive isolation is established [1-3]. According to the genic view of spéciation [4], when populations are in contact, gene flow is initially restricted at barrier loci (i.e. loci underlying reproductive isolation), leading to the emergence of peaks of genetic differentiation surrounding such barriers; i.e. heterogeneous genomic differentiation [5,6]. As spéciation progresses, this localised build-up of reproductive isolation spreads to nearby regions due to linkage [4,5,7], Once a critical amount of differentiation at multiple barrier loci has accumulated, reduction of the genome-wide effective migration rate will eventually lead to divergence across the entire genome [5,7], This final step of genome-wide congealing may be a rapid and non-linear phase transition under certain conditions, such as when isolating barriers have a polygenic basis or a few strong barrier loci arise [8-10]

Recent empirical genomic studies have revealed regions of high and low differentiation dispersed throughout the genome at early stages of spéciation This empirical data has lent strong support to the genic perspective of the spéciation process [4], To-date however, the majority of spéciation genomic studies demonstrating heterogeneous genetic differentiation have come from young species or population pairs with low divergence [7,11,12] with some exceptions [9,13]. Even for these cases [13,14], it is unclear whether gene flow has occurred throughout divergence or whether the species pair in question experienced periods of geographical isolation (see below). A bias towards early stage divergence has undoubtedly been informative to understand the onset of the spéciation process [15-18]. However, it is limited in its scope for understanding the factors that eventually lead to the completion of the spéciation process - i.e. the evolution of genome-wide differentiation. When divergence with gene flow is studied across the spéciation continuum in *Timema* stick insects (including the later stage divergence) the range of genome-wide differentiation appears to be disjointed with a gap in *F_st_* between 0.3 and 0.7 [8]. Furthermore, literature surveys have also indicated that many species pairs exhibit a mean *F_st_* between either 0.03-0.27 or 0.70-0.90 when gene flow is occurring [8,9]. This disjointed distribution of differentiation is consistent with the idea that progression towards spéciation may be non-linear, with a phase transition due to genome wide congealing [10,19].

In addition to need for more balanced understanding of the extent of genomic divergence and introgression at later stages of the speciation continuum, there is a need for studies which account for factors such as demographic history [12,14], This is because high genome-wide differentiation in a single species pair may have evolved via genetic drift and local adaptation during allopatric isolation, rather than due to divergence with gene flow. In other cases, heterogeneous genomic differentiation may be due to erosion of genetic differentiation due to introgression following secondary contact after geographical isolation. Without a picture of the demographic history, this scenario may be indistinguishable from primary divergence [20] and will therefore introduce bias if multiple species pairs are compared. Despite the fact that the expected pattern of genomic differentiation during speciation is influenced by the timing and duration of geographical isolation [7], testing different demographic histories has been somewhat neglected by the field [7,20].

Other factors besides demographic history of a species pair can also confound patterns of heterogeneous genomic differentiation. For example, local adaptation or background selection in genomic regions where recombination is reduced can elevate differentiation measures and be mistaken for barrier loci [13,21,22], Similarly, regions of low differentiation may be caused by not only on-going gene flow but also incomplete lineage sorting (ILS) [21,23] and mutation rate variation [24], Distinction between gene flow and ILS is likely easier in more divergent species pairs [24-26]. Furthermore, the use of multiple classical and recently developed methods, such as detection of recent hybrid progeny, ABBA-BABA tests [27,28], model-based inference[29], and comparison between allopatric and sympatric pairs [22,28] provide a means to distinguish signatures of gene flow from alternative explanations. Thus, there is a need for studies which account for factors such as demographic history, recombination rate variation, and ILS that can confound the interpretation of genome scan data [7,12].

Three-spined stickleback species pairs (genus *Gasterosteus)* span the speciation continuum at varying stages of divergence, making them a model system for speciation research [30,31]. To-date genomic research on speciation with gene flow in the stickleback complex has largely focused on weakly divergent species pairs, such as lake-stream ecotypes, with mean genome-wide *F_st_* values of less than 0.3 [32-34], Such studies have shown that the genomic landscape of differentiation between these recently diverged sympatric or parapatric species pairs is heterogeneous and interspersed with multiple peaks of high differentiation [16,32,34], The emerging pattern is consistent with predictions under the genic concept of speciation - i.e. that reproductive isolation is localized in the genome at early stages of speciation [4,35]. However, it remains unclear whether such localized differentiation will eventually progress toward genome-wide differentiation in the face of gene flow.

Toward the end of the stickleback speciation continuum is a marine species pair in Japan [36,37], The Japan Sea stickleback (*G. nipponicus*) is sympatric with the Pacific Ocean lineage of three-spined stickleback (*G. aculeatus*) (Fig 1A) in the waters surrounding the Japanese archipelago (Fig 1C) [36,38], Divergence time between the two marine species has been estimated to be 1.5-2 million years based on allozyme and microsatellite data [37,39], making it much older than postglacial stickleback species pairs. Speciation has been hypothesized to have occurred as a result of the repeated isolation of the Sea of Japan during the Pleistocene, but this divergence scenario remains to be explicitly tested [37,39]. A unique feature of the *G. nipponicus* and *G. aculeatus* system, relative to postglacial stickleback species pairs, is that a neo-sex chromosome has arisen due to a fusion between a Y chromosome and a previously autosomal chromosome IX (chrIX) in the *G. nipponicus* lineage [36,40]. Furthermore, crosses between Japan Sea females and Pacific Ocean males show hybrid male sterility [37], Previous quantitative trait locus (QTL) mapping identified QTL for courtship behaviour on the neo-X and hybrid male sterility on the ancestral-X. However, there are other isolating barriers, such as eco-geographical isolation, temporal isolation, and ecological selection against migrants [37,41,42], The combination of these multiple barriers most likely contributes to the strong reproductive isolation in this system [36,43], However, despite such strong divergence, hybrids have been observed where the two species co-occur in Northern Japan [36] and mitochondrial discordance between the species suggests some history of introgression during speciation [44,45], Although the Japanese species pair therefore represents the furthest point of divergence along the stickleback speciation continuum, speciation remains incomplete. The evolutionary history and genome-wide patterns of genetic differentiation and introgression of this strongly divergent species pair therefore remains an open question

**Fig 1.**
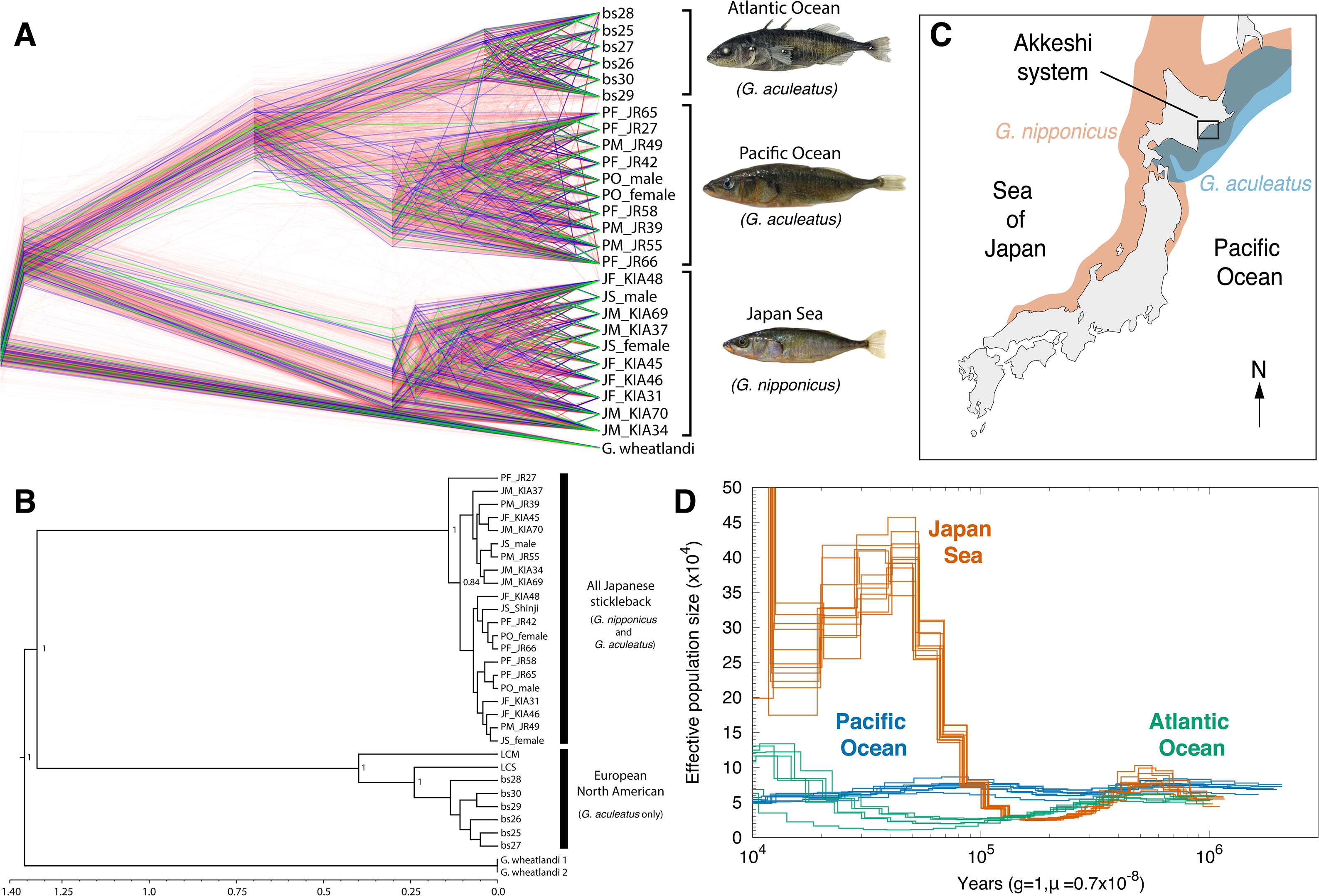
The Japan Sea stickleback is a separate species. (A) Rooted nuclear consensus tree for Japan Sea, Pacific Ocean and Atlantic Ocean stickleback lineages from 10 kb non-overlapping sliding windows across the autosome. Red trees indicate species clustering; blue trees indicate geographical clustering and green trees reflect ancestral polymorphism. NB: Only 1,000 subsampled species trees are shown here to aid illustration. (B) Mitogenome Bayesian consensus tree shows divergence between two mitochondrial clades - all Japanese sticklebacks (*G. nipponicus* and *G. aculeatus*) and *G. aculeatus* occurring in Europe and North America. (C) Present day distribution of *G. aculeatus* (blue) and *G. nipponicus* (red) around the Japanese archipelago. The two species overlap in Hokkaido, Northern Japan and samples for this study were collected in Bekanbeushi River in Akkeshi unless noted. (D) PSMC plot of 26 resequenced genomes shows a steady population size in the Pacific Ocean lineage (blue) but a bottleneck around 0.15-0.3 million years before present and a subsequent increase in the Japan Sea lineage (orange).

The aim of our study was to address gap in our knowledge; i.e. to quantify the patterns of genomic differentiation and introgression at a later stage of the speciation continuum. To this end, we used a whole-genome reseqeuncing data from the Japanese stickleback species pair to determine their evolutionary history and characterise patterns of gene flow between them. Our first aim was to establish how and when divergence took place between *G. nipponicus* and *G. aculeatus*. Using thousands of genomic loci and a coalescent modelling approach, we tested a range of divergence scenarios and estimated the timing and duration of isolation, the extent of gene flow and fluctuations in population size. After identifying that the two species have indeed diverged in the face of gene flow, we then used a comparative genome scan approach with an additional *G. aculeatus* lineage from the Atlantic Ocean [46] as an allopatric control (Fig 1A, Fig SI). After establishing that gene flow has occurred but that a high level of genomic differentiation has remained, we used two independent measures of gene flow to identify where in the genome introgression has left its mark. We tested whether introgression occurs more frequently in regions of high recombination and whether it occurs in regions with functionally important genes. Our findings suggest a high level of genome-wide divergence can be maintained, as introgression is restricted to small, localized genomic regions.

## Results

### Ancestral demography and population genomic analyses support divergence with gene flow

Phylogenetic analysis on 35,666 10 kb non-overlapping genome windows on autosomes (i.e., excluding chrIX and chrXIX) supports a deep split between *G. aculeatus* (both Pacific and Atlantic Ocean lineages) and *G. nipponicus* (Japan Sea stickleback) (Fig 1A). Of all windows, 98.8% support the split between species, while only 0.51% indicate clustering of fish occurring in Japan (the Japanese Pacific Ocean *G. aculeatus* and the Japan Sea *G. nipponicus;* Table SI and Fig 1A).

We calculated genealogical sorting index *(gsi*) [47] on maximum likelihood phylogenies estimated from non-overlapping sliding windows of 20 kb across the autosome. High *gsi* indicates monophyly, while low *gsi* indicates mixed ancestry [47], Genome-wide averages (± SD) of *gsi* were high, but not complete, for all three *Gasterosteus* lineages with that of the Japan Sea stickleback being the highest (Atlantic *gsi =* 0.45 ± 0.10, Pacific *gsi =* 0.57 ± 0.09, Japan Sea *gsi =* 0.72 ± 0.06).

This is in stark contrast to the mitogenome phylogeny where sticklebacks from both species occurring in Japan fall into a single clade separate from the clade occurring in the Western Pacific and Atlantic (Fig IB, Fig S2). A lack of mitogenome divergence between *G. aculeatus* and *G. nipponicus* from the Japanese archipelago suggests mitochondrial introgression has occurred where these lineages overlap (Fig 1C).

Since the consensus autosomal phylogeny suggests a more recent split between the Pacific and Atlantic *G. aculeatus* lineages, a deeper mtDNA split based on spatial distribution suggests mitochondrial introgression has likely occurred from the Japan Sea *G. nipponicus* into the Pacific Ocean *G. aculeatus*. Divergence time estimates between the mitogenome clades are thus informative for dating speciation. Bayesian coalescent analysis using a strict clock model in Bayesian Evolutionary Analysis by Sampling Trees (BEAST) suggests a median split date of 1.30 million years (0.15-2.41; 95% Highest Posterior Density [HPD] intervals; Table S2) for the two major mitogenome clades (Fig S2), consistent with previous estimates [44], Divergence between Eastern Pacific and Atlantic haplotypes is more recent at 0.39 million years (0.03-0.74; 95% HPD) but is older than the Most Recent Common Ancestor (MRCA) of all haplotypes occurring in Japan (Fig IB, S2A), suggesting mitochondrial gene flow from *G. nipponicus* to *G. aculeatus* has occurred in the recent past (i.e. <0.39 million years BP).

Mitochondrial introgression can be driven by large demographic disparities between populations with gene flow and is more likely to occur from a larger to a smaller population [48]. To investigate the demographic disparities between *G. aculeatus* and *G. nipponicus*, we used pairwise sequential Markov coalescent (PSMC) on all 26 Atlantic Ocean, Japan Sea and Pacific Ocean resequenced stickleback genomes. Strikingly, *G. nipponicus* experienced a severe bottleneck around 0.15-0.3 million years before present (BP) (Fig ID); mean *N*_*e*_ fell to 26,422 ± 1,191 at its lowest point. Subsequently after 0.1 million years BP, *G. nipponicus* underwent a dramatic population size expansion (Fig ID): mean *N*_*e*_ rose to 195,974 ± 28,832 (i.e.∼7.5 times increase from the bottleneck) during the late Pleistocene. In contrast, the Japanese Pacific Ocean *G. aculeatus* population has remained relatively stable throughout its history (mean *N*_*e*_± SD = 118,150 ± 4,330; Fig ID, see Fig S3 for bootstrap support). Although the Atlantic (Fig ID) and Western Pacific lineages of *G. aculeatus* (Fig S4) also experienced some population growth during the late Pleistocene, their effective population sizes remained smaller than that of *G. nipponicus*. Genome-wide averages of Tajima’s *D* also support a recent demographic expansion for *G. nipponicus* (mean ± SD of Tajima’s *D =* -0.82±0.45) and stable population size in the Pacific Ocean (mean ± SD of Tajima’s *D =* -0.04 ± 0.63). Taken together, these data indicate that mitochondrial introgression likely occurred during the late Pleistocene, when *G. nipponicus N_e_* was substantially larger than *G. aculeatus N_e_*. This is consistent with the hypothesis that mitochondrial gene flow from *G. nipponicus* to *G. aculeatus* has occurred <0.39 million years BP (see above).

To explicitly test whether divergence between *G. aculeatus* and *G. nipponicus* occurred in the presence of gene flow, we used an Approximate Bayesian Computation (ABC) approach with 1,874 2 kb loci randomly sampled from across the autosome. We tested five divergence scenarios - isolation (I), isolation with migration (IM), isolation-with-ancient-migration (IAM), isolation-with-recent-migration (IRM) and isolation-with-ancient-and-recent-migration (IARM) - i.e. two discrete periods of contact Since the results of our PSMC analyses clearly indicate *N*_*e*_ has varied throughout divergence (Fig ID), we performed a hierarchical ABC analysis, first selecting the most appropriate population growth model (i.e. constant size, population growth and a Japan Sea bottleneck) within each divergence scenario and then performing final model selection amongst the best supported divergence/growth model scenarios (see Supplementary Methods for full specification of models, priors, parameters and extensive sensitivity testing).

Using 20 summary statistics (see Supplementary Methods for a full list of statistics used) and a neural-network rejection method with 1% tolerance of simulated datasets, the best-supported divergence scenario was a model of IM (Fig 2A; Table 1). Parameter estimates from the IM model suggest divergence between *G. aculeatus* and *G. nipponicus* occurred 0.68 million years ago (median estimate, 0.18-4.17 million years, lower & upper 95% HPD; Fig 2B). A Japan Sea bottleneck occurred 0.3 million years ago (0.03-2.21 million years 95% HPD), reducing *N*_*e*_ to about 20% of the contemporary estimate (Fig 2C, Table S3). Mean migration rates between the two species were low, and migration from the Japan Sea into the Pacific Ocean lineage was slightly greater (Fig 2D, Table S3). Contemporary *N*_*e*_ of the Japan Sea lineage is larger than that of the Pacific Ocean, although the *N*_*e*_ estimates differed in magnitude from those estimated by PSMC (Figs ID and 2C, Table S3).

**Table 1.**
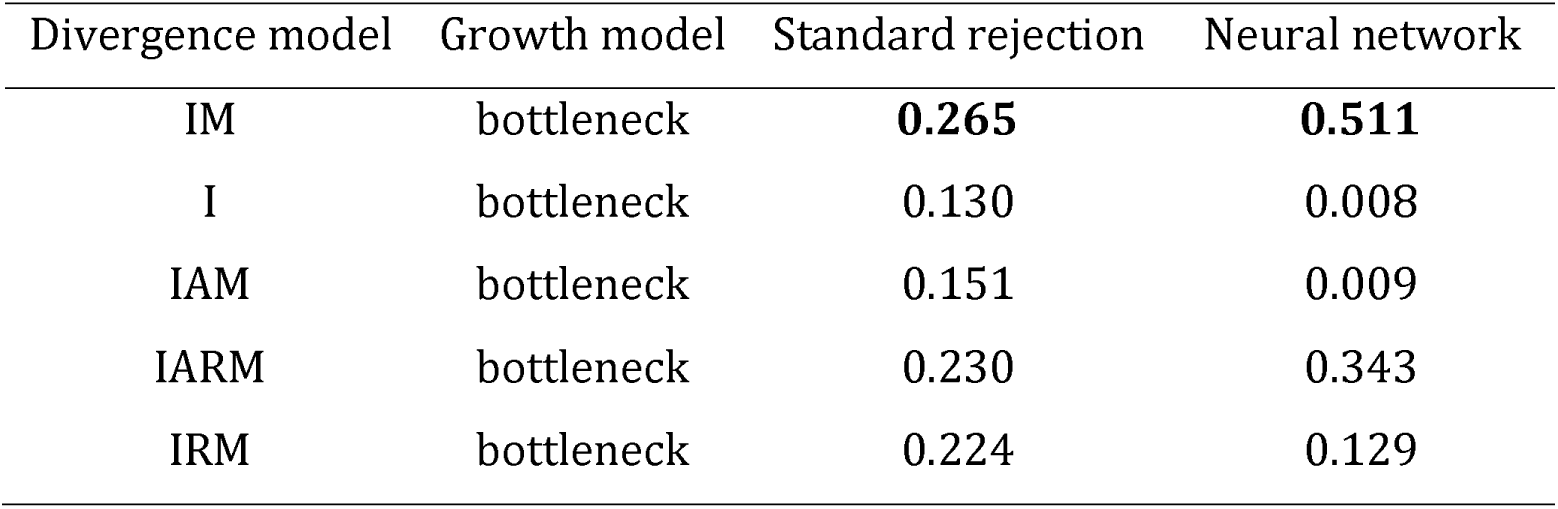
Posterior probability values for models for final ABC model selection using neural network and standard rejection methods. All estimates produced using a tolerance of 1% and 20 summary statistics. Bold text indicates the model where posterior probability provides the highest support Models are I = isolation, IM = isolation with migration, IAM = isolation and ancient migration, IRM = isolation and recent migration, IARM = isolation with ancient and recent migration.

**Fig 2.**
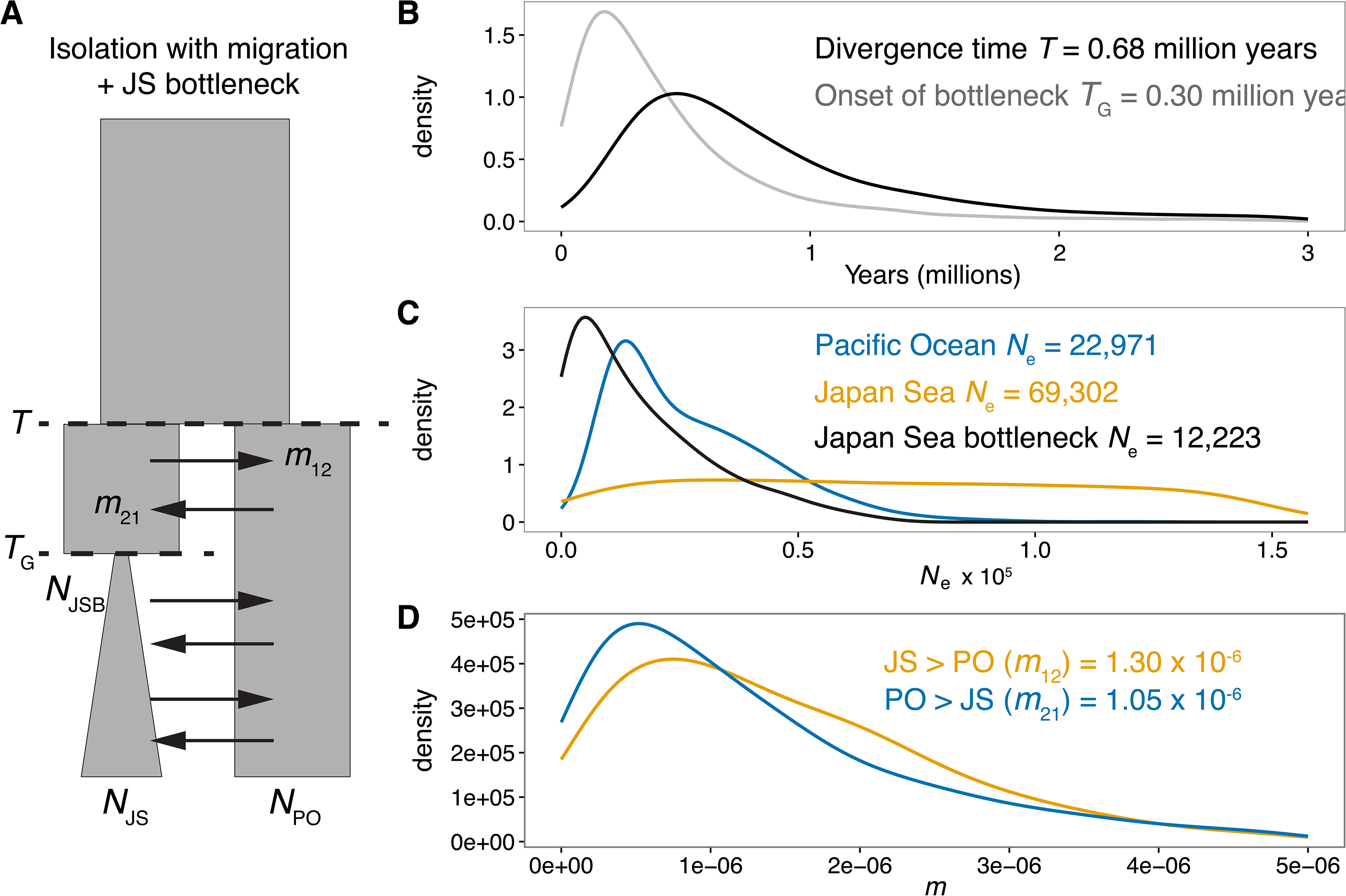
ABC analysis supports isolation with gene flow. (A) A model of isolation with migration and a bottleneck in the Japan Sea lineage is best supported by ABC analysis using —2,000 nuclear loci (see Table 1). Posterior probability densities for model parameters estimated using neural network analysis, a 1% tolerance rate and 20 summary statistics. Parameters are: T = time of split, *m*_12_ *=* migration from Japan Sea to Pacific Ocean, *m*_21_ = migration from Pacific Ocean to Japan Sea; *T*_G_ *=* timing of bottleneck, *N*_PO_ = Pacific Ocean effective population size, *N*_JS_ = Japan Sea effective population size and *N*_JSB_= Japan Sea bottleneck effective population size. Posterior probability density curves for (B) Japan Sea and Pacific Ocean divergence time and timing of bottleneck in the Japan Sea lineage, (C) Japan Sea, Pacific Ocean and Japan Sea bottleneck effective population size. Posterior probability density curves for (B) Japan Sea and Pacific Ocean divergence time and timing of bottleneck in the Japan Sea lineage, (C) Japan Sea, Pacific Ocean and Japan Sea bottleneck effective population sizes, and (D) mean migration rates. Figures on each panel are median parameter estimates.

Identifying admixture between species where they co-occur provides strong evidence of on-going introgression [7,12], To address this, we used a RAD-sequencing dataset with a larger sample size of 245 individuals from the Atlantic, Pacific and Japan Sea lineages, including previously published data from Pacific-derived populations in North America [49]. Principal component analysis (PCA) of allele frequencies at 3, 744 high-quality bi-allelic SNPs showed that, consistent with our whole genome data, the main axis explaining 20% of the variance was between *G. aculeatus* and *G. nipponicus* (Fig S5). The secondary axis explaining 9.49% of the variance was mainly between the Atlantic and Pacific populations (Fig S5). Importantly, PCA shows a single individual is intermediate between the Pacific and Japan Sea populations occurring in Akkeshi, the sympatric site in Hokkaido, Japan where our whole genome-sequenced samples were collected (Fig 1C). A separate Bayesian analysis for admixture using STRUCTURE [50,51] found greatest support for *K =* 2 in the Japanese populations and also identified the putative F_1_ hybrid plus individuals with possible recent admixture at this sympatric site (Fig S6).

Taken together, these data indicate that divergence between the Japanese *G. aculeatus* and *G. nipponicus* is much older and greater compared to commonly studied postglacial stickleback species pairs. Despite the great extent of divergence between Japanese stickleback species, parameter estimates and observational data suggest that gene flow between them is on-going.

### High levels of genome-wide divergence with highly localized signatures of introgression

Genome-wide differentiation was strikingly high between *G. nipponicus* and *G. aculeatus* regardless of their geographical overlap: both relative (*F*_ST_) and absolute divergence (*d*_XY_) were high (Fig 3A & B, Fig 4, and Figs S7 and S8). The genome-wide average of *F*_ST_ between the sympatric species was 0.628; this is higher than all other studied stickleback species pairs [32-34,52] (see Fig 3C). Despite consistently high divergence, both *F*_ST_ and *d*_XY_ values were significantly lower where the two species occur in contact (Table 2, Figs 3A & B, 4, S7 and S8; 10,000 replicate permutation tests on 10 kb windows: *P < 2.2 ×* 10^16^ for both statistics), consistent with the presence of gene flow in sympatry.

**Table 2.** Genome-wide averages for measures of divergence and introgression. *F*_ST_, *d*_XY_, *G*_MIN_, and *f*_d_ for all pairwise comparisons of Japan Sea (JS), Pacific Ocean (PO) and Atlantic Ocean sticklebacks (AT) are shown. Mean ± SD and lower and upper limits of the 95% confidence interval (in parenthesis) are shown. NA, not analysed.

| Comparison | $F_{ST}$ | $d_{XY}$ | $G_{MIN}$ | $f_d$ |
| --- | --- | --- | --- | --- |
| JS vs PO | $0.634 \pm 0.122$ | $0.012 \pm 0.002$ | $0.857 \pm 0.102$ | $0.004 \pm 0.054$ |
|  | (0.333-0.862) | (0.007-0.017) | (0.513-0.942) | (-0.031-0.085) |
| JS vs AT | $0.697 \pm 0.116$ | $0.013 \pm 0.002$ | $0.876 \pm 0.071$ | $-0.003 \pm 0.033$ |
|  | (0.406-0.902) | (0.007-0.018) | (0.666-0.942) | (-0.077-0.029) |
| PO vs AT | $0.215 \pm 0.134$ | $0.004 \pm 0.002$ | $0.560 \pm 0.141$ | NA |
|  | (0.003-0.539) | (0.001-0.009) | (0.223-0.772) |  |

**Fig 3.**
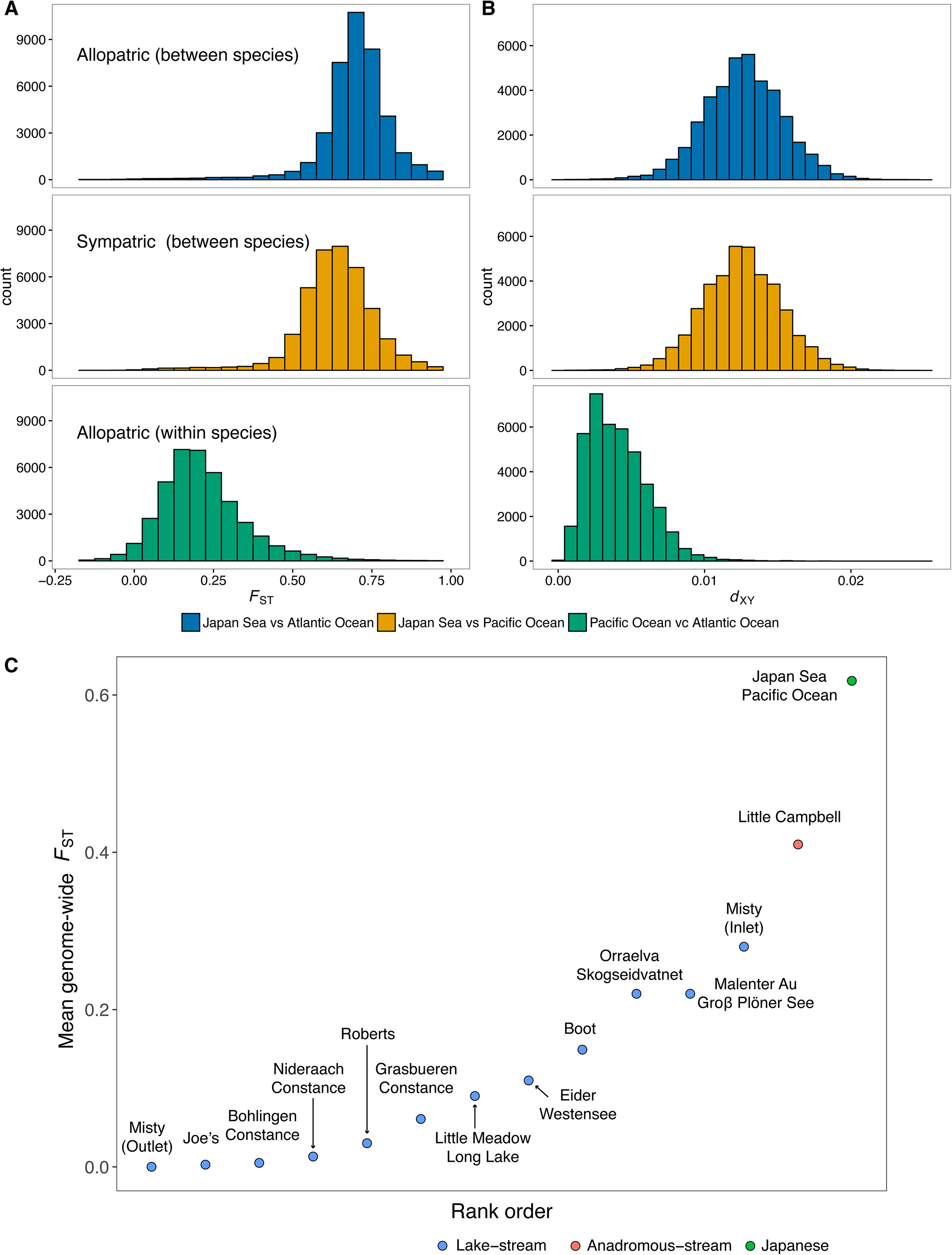
Genomic divergence is lower in sympatry than in allopatry between species. Histograms of [A] relative (*F*_ST_) and (B) absolute (*d*_XY_) differentiation measures for each of the species comparisons. (C) Mean genome-wide *F*_ST_ of the Japanese species pair compared with those of other stickleback systems taken from previously published studies [32-34,52].

**Fig 4.**
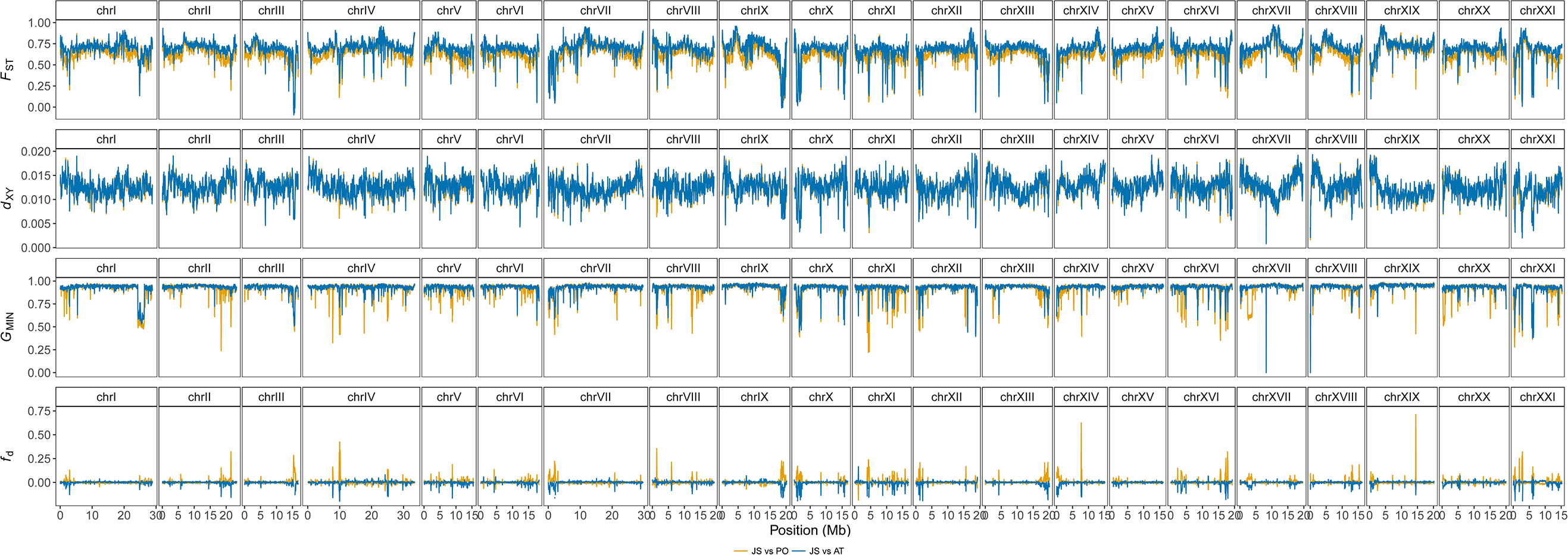
Genome-wide distribution of divergence and introgression. Divergence was measured using *F*_ST_ and *d*_XY,_ while introgression was measured using G_MIN_ and *f*_d_. Data plotted here is from 50 kb non-overlapping genome windows. Blue and yellow lines indicates allopatric (Japan Sea vs Atlantic) and sympatric (Japan Sea vs Pacific Ocean) comparisons, respectively.

A more fine-scale analysis of genome-wide divergence based on 10 kb non-overlapping windows revealed that the high baseline divergence between *G. nipponicus* and *G. aculeatus* is interspersed by regions of low differentiation in both *F*_ST_ and *d*_XY_ genome scans (Figs 4 top two panels, S7 and S8). Processes such as background selection that alter within-population diversity can bias relative measures of differentiation such as *F*_ST_ [7,21,22,53], Furthermore, absolute measures like *d*_XY_ can take a long time to reach equilibrium between populations following divergence, meaning a lower power for detecting recent introgression events [7,26]. Therefore, to better identify genomic regions of recent introgression, we calculated two independent measures of introgression. The first of these was *G*_MIN_, the ratio of the minimum *d*_XY_ to the average *d*_XY_ [26]. Under strict isolation, *G*_MIN_ is the lower bound of divergence time between two populations, whereas when introgression occurs, *G*_MIN_ reflects the timing of the most recent migration [26]. The second measure was *f*_d_, an estimate of the proportion of introgressed sites in a genome window, calculated using a four population ABBA-BABA test [54], *G*_MIN_ is more effective at identifying recent, low level gene flow than either *F*_ST_ or *d*_XY_ but by definition it is unable to detect genomic regions where complete introgression has occurred [26], which can however be detected using *f*_d_. Importantly, both measures are robust to variation in recombination rate [26,54], Combining these two statistics therefore allows us to identify both low-level (*G*_MIN_) and strong introgression (*f*_d_).

Focusing on between species comparisons, mean (± SD) *G*_MIN_ measured from 10 kb non-overlapping windows was greater in allopatry than sympatry (Japan Sea vs. Atlantic: 0.876 ± 0.071; Japan Sea vs. Pacific: 0.857±0.103; randomization testP < 2.2xl0^-16^; Fig 4). Mean/d was also greater when the species overlapped (JS vs. AT: -0.0031 ± 0.0540; JS vs. PO: 0.0039±0.0328; *P* < 2.2xl0^-16^; Fig 4), and both statistics are more strongly negatively correlated in sympatry (Fig S9) supporting gene flow between *G. nipponicus* and Japanese populations of *G. aculeatus*.

Genomic regions of low *G*_MIN_ (i.e. *G*_MIN_ valleys) may indicate recent introgression. We identified genome windows with low *G*_MIN_ values using a Hidden-Markov classification model [55] and an outlier approach based on permutations of variant sites (Fig S10). We then clustered 10 kb outlier windows occurring within 30 kb of one another into putative *G*_MIN_ valleys. *G*_MIN_ in particular may be susceptible to false positives as a result of ILS. However, lower *d*_XY_ and higher *f*_d_ in sympatric *G*_MIN_ valley windows compared to the genomic background suggests ILS alone does not explain the patterns observed here (Fig S11; randomization test, *P < 2.2 ×* 10^-16^in both cases). These regions of most recent migration [26]. The second measure was *f*_d_, an estimate of the proportion of introgressed sites in a genome window, calculated using a four population ABBA-BABA test [54], *G*_MIN_ is more effective at identifying recent, low level gene flow than either *F*_ST_ or *d*_XY_ but by definition it is unable to detect genomic regions where complete introgression has occurred [26], which can however be detected using *f*_d_. Importantly, both measures are robust to variation in recombination rate [26,54], Combining these two statistics therefore allows us to identify both low-level (*G*_MIN_) and strong introgression (*f*_d_).

A similar geographical comparison of peaks of *f*_d_ between species was not possible, due to the much lower genome-wide distribution of *f*_d_ between *G. nipponicus* and the Atlantic *G. aculeatus* (Fig 4). Nonetheless, Hidden-Markov classification identified 823 *f*_d_ peaks occurring between *G. nipponicus* and Pacific *G. aculeatus* (Fig S12). If the *f*_d_ peaks are mainly indicate introgression from Pacific Ocean to Japan Sea, *d*_XY_ between Japan Sea and Atlantic Ocean is expected to be lower in these regions compared to the genome background, as Japan Sea fish carry haplotypes derived from the Pacific Ocean, which in turn are similar to the Atlantic Ocean haplotypes. While JS-AT *d*_XY_ was lower in *f*_d_ peaks compared to the genome background (one-tailed permutation test, *P* < 2.2 × 10^16^), this difference was not very clear (Fig. S13). In contrast, if introgression occurred mainly from Japan Sea to Pacific Ocean, dxyin the PO-AT comparison should increase in *f*_d_ peaks relative to the genome background, as Pacific Ocean fish carry Japan Sea-derived haplotypes, which are divergent from the Atlantic Ocean haplotypes. We clearly observed this pattern (P < 2.2 × 10^-16^; Fig. S13); suggesting that introgression from Japan Sea to Pacific Ocean may be more predominant than the opposite direction.

To further investigate the direction of gene flow, we used partitioned *D*) statistics (an extension of the four population test - see Fig. S14), which tests the excess of shared derived alleles using five, rather than four populations [56]. To this end, we added an allopatric Japan Sea population (collected from Lake Shinji, a brackish lake at the Japan Sea coast of southern Honshu). A positive D_12_ statistic is proposed to indicate the predominance of introgression from P3 to P2 (Fig. S14) [56]. When P3 was set to Japan Sea (where P3_1_ is sympatric and P3_2_ is allopatric with the Pacific Ocean) and P2 to Pacific Ocean (see Fig. S14B), *D*_*i2*_ was significantly positive in *f*_d_ peaks (one-tailed permutation test, P < 2.2 × 10^-16^). In contrast, when we rotated the populations at the tips - i.e. setting P2 to sympatric Japan Sea, P3_1_ to Pacific Ocean, and P3_2_ to Atlantic Ocean (see Fig S14C), D12 was not positive, consistent with the suggestion that introgression is occurring mainly from Japan Sea to Pacific Ocean. However, the resolution of partitioned *D* statistics has been criticized [57]; positive D_12_ can also be caused by introgression from the Pacific Ocean (P2) to the common ancestor of the sympatric and allopatric Japan Sea populations (P3_1_ & P32). To overcome this issue, we calculated Dfoil, which also uses a five-population test but accounts for all possible introgression events [57], When P_1_= sympatric Japan Sea, P2 = allopatric Japan Sea, P3 = Pacific Ocean, and P4 = Atlantic Ocean (Fig. S15), Dfoil clearly indicated the presence of ancestral introgression (239 out of 4,236 100 kb-windows) between the Japan Sea ancestor (P12) and the Pacific Ocean (P3) (see Fig. SI5). However, we found only a few windows showing unidirectional introgression (6 loci in total), precluding conclusion of the predominant direction of introgression using this analysis (Fig. S15). This low sensitivity may be due to the fact that structuring in the Japan Sea lineage is low [58] - i.e. recent divergence time or high intraspecific gene flow.

### Characterization of genomic regions of introgression

To investigate whether introgression co-varies with recombination rate, we used a previously published recombination map from an Atlantic *G. aculeatus* cross [59] to interpolate genome-wide recombination rate variation (see Methods). We detected a negative correlation between recombination rate and Cm in and a positive correlation with/d (Pearson’s correlation, *G*_MIN_: r = -0.17, *P < 2.2* × 10^-16^; *f*_d_: r = -0.08, *P< 2.2 ×* 10^-16^, Fig S16). Accordingly, mean recombination rate for putatively introgressed regions was over two times higher than the genome background (*G*_MIN_: valley = 8.98 cM/Mb, non-valley = 3.99 cM/Mb;*f*_d_: peak = 9.64 cM/Mb, non-peak = 4.16 cM/Mb; randomization testP < *2.2 ×* 10^16^in both cases; Fig 5B).

**Fig 5.**
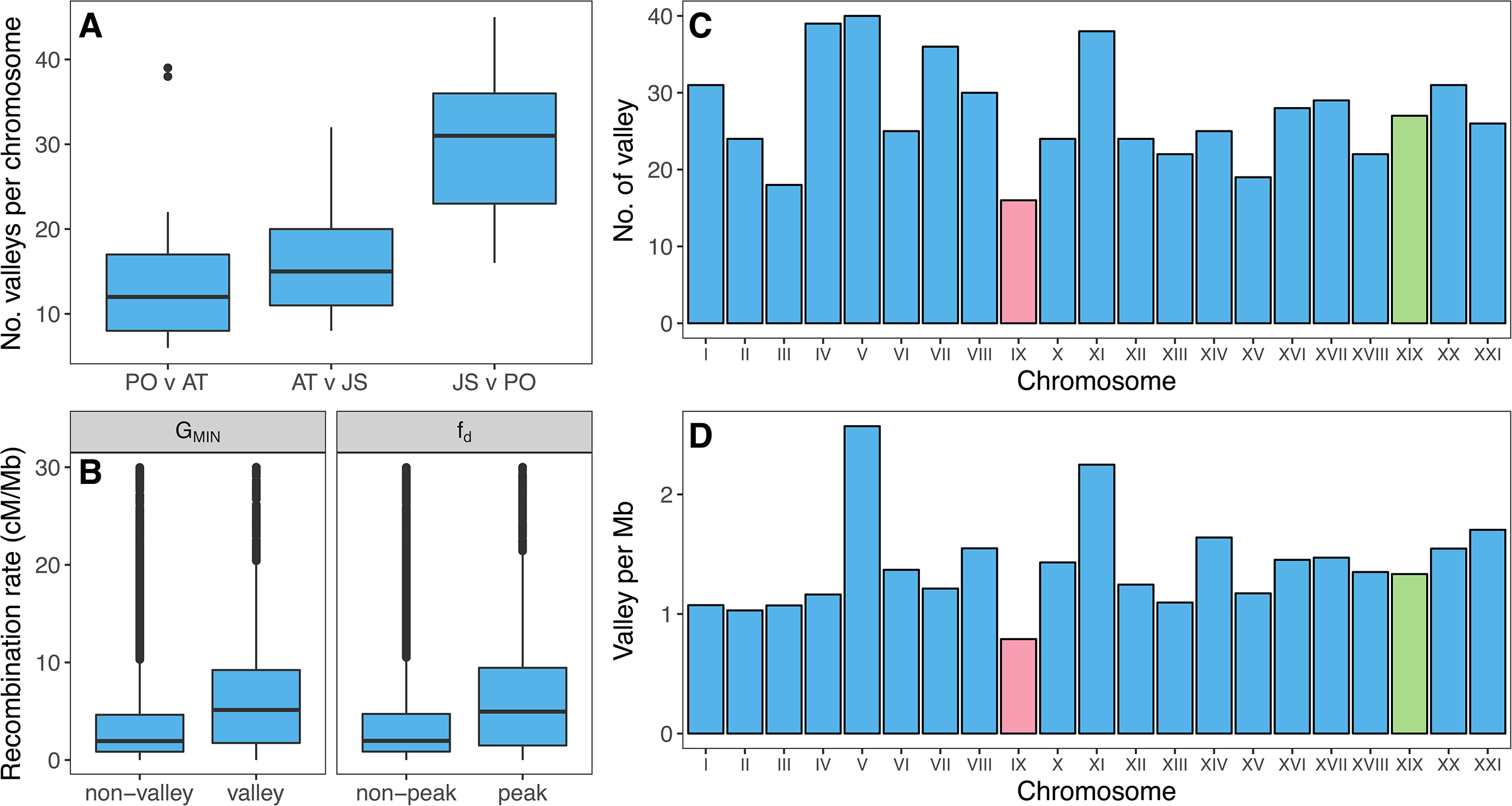
Fewer introgression valleys occur on the neo-X chromosome. (A) A greater number of *G*_MIN_ valleys occur in sympatry than in allopatry between species. (B) *G*_MIN_ valleys and *f*_d_ peaks also occur in regions of the genome with a higher recombination rate. Fewer valleys occur on the neo-X chromosome (chrIX) compared to autosomes (C), even when chromosome length is taken into consideration (D); N.B - data for (C) and (D) were measured using females only.

Sex chromosomes likely played an important role in speciation between *G. aculeatus* and *G. nipponicus*. A fusion between Y and chrIX means that chrIX segregates as a neo-sex chromosome in *G. nipponicus* but not *G. aculeatus* which only carries the ancestral and shared sex chromosome, chrXIX [36,40]. The divergent XY *(G. aculeatus*) and X1X2Y *(G. nipponicus*) systems means that recombination is reduced for chrIX and chrXIX in hybrids carrying the neo-Y [40]. Given this recombination rate reduction and previously identified QTL for traits involved in reproductive isolation that map to chrIX and chrXIX [36,40], we tested whether recent introgression (i.e. measured using *G*_MIN_) was reduced in this part of the genome relative to the autosome. For this, we repeated our analyses using females only (5 Japan Sea and 6 Pacific Ocean). The number and density of valleys was lowest on the neo-sex chromosome, chrIX (16 valleys or 0.8 valleys per Mb) but not the ancestral sex chromosome (chrXIX, see Table S4).

Finally, we investigated the nature of introgression between the two species. We first asked whether introgression occurs more frequently in genic or non-genic regions. We identified 3,261 genes occurring in *G*_MIN_ valleys and 2,958 genes from *f*_d_ peaks between sympatric *G. aculeatus* and *G. nipponicus;* 60% of genes identified were found in both types of introgressed window, whereas 23% occurred only in Gmin valleys and 15% only in *f*_d_ *peaks* (see Fig S17). Irrespective of the method used to detect putatively introgressed regions, the number of genes identified was greater than the number expected by chance (P < 0.0001 based on a null distribution generated from 1,000 random samples of the genome). This suggests that introgression may be more likely in genic regions of the genome than non-genic regions.

To further investigate the functional enrichment of the genes occurring in regions of introgression, we performed gene ontology (GO) analysis on 2,310 *G*_MIN_ valley and 2,217 *f*_d_ peak genes with orthologs in the human genome. Enriched GO terms for *f*_d_ peaks included immune response, metabolic processes and chromatin assembly, while enriched GO terms for *G*_MIN_ valleys included major histocompatibility complex (MHC) protein and metabolic processes (Table S5&S6).

## Discussion

### Japanese stickleback spéciation has occurred in the face of on-going gene flow

Determining the demographic and evolutionary history of species pairs is an important first step for understanding how spéciation has unfolded in any system [7, 12]. Our present study has produced several lines of evidence indicating that divergence between the Japanese sticklebacks has occurred in the presence of gene flow.

Firstly, contrasting mitochondrial and nuclear genome phylogenies show that mitochondrial introgression has occurred from *G. nipponicus* into *G. aculeatus* at some point in the last 0.39 million years. Our mitogenome phylogeny confirmed previous findings that there is no mitochondrial divergence between the *G. nipponicus* and Japanese populations of *G. aculeatus* [44,45]. This is in contrast to our nuclear autosomal phylogeny which showed that majority of the genome supports a clear split between *G. nipponicus* and *G. aculeatus* occurring in Japan and that the latter shares a more recent common ancestor with Atlantic European *G. aculeatus* populations. In short, mitogenome data clusters the *Gasterosteus* lineages by geography, while the nuclear data clusters them by species. Introgression is the most likely explanation for this mitonuclear discordance as the lower effective population size of the mitochondrial genome relative to the autosome makes ILS unlikely, particularly over a 1 million year time-scale of divergence.

Disparities in effective population size between lineages are a common cause of unidirectional mitonuclear introgression with introgression likely occurring from a larger to a smaller population [48]. Our reconstruction of temporal variation in effective population size using PSMC showed a rapid population explosion *ofG. nipponicus* during the late Pleistocene that created a large demographic disparity with the *G. aculeatus* Pacific Ocean lineage, although it should be noted that admixture can increase effective population size estimates when using PSMC [60,61]. Reasons for this population growth remain unclear but it is surprising, particularly since *G. nipponicus* is unable to colonise freshwater environments [42,58], which might be expected to increase effective population size due to meta-population dynamics. A possible explanation is that a lack of dependence on freshwater for spawning [41,62,63] and greater foraging efficacy on marine prey [42] means *G. nipponicus* is better adapted for exploiting an abundant marine environment Unidirectional mitochondrial introgression might also be caused by female mate choice [64], Our previous behavioural studies indicate that Japan Sea females often mate with Pacific Ocean males, while Pacific Ocean females rarely mate with Japan Sea males [36,37], Hybrid females from Japan Sea female and Pacific Ocean male crosses are fertile [37] and will carry Japan Sea mitochondrial DNA. Backcrossing of these hybrids to Pacific Ocean males would result in unidirectional mitochondrial introgression from the Japan Sea to Pacific Ocean.

Secondly, our ABC analysis supported a model of isolation with migration. Previously, it has been speculated that the Japan Sea stickleback diverged largely as a result of geographical isolation in the Sea of Japan caused by sea level fluctuation during the early Pleistocene [37,39]. Using ABC, we were able to explicitly test several divergence hypotheses in a statistical framework [65]; our findings suggest that gene flow has likely occurred throughout the majority of divergence history. It should be noted that ABC and most established demographic inference methods perform poorly when resolving the timing of gene flow between lineages [66,67], Therefore, one caveat to the interpretation of our ABC results is that we cannot rule out the possibility that the two species diverged in repeated cycles of contact (i.e. akin to our IARM model which had the second highest level of support; Table 1), but these periods of contact were simply too close in time. Nonetheless, the pooled posterior probabilities from the analysis overwhelmingly support a model of divergence with gene flow irrespective of the timing or nature of the actual speciation event The presence of extant recent hybrids in sympatry also strongly indicates that hybridization is still on-going. We observed a probable F_1_ hybrid in the wild and several other individuals with evidence of hybrid ancestry in our RAD-seq dataset, consistent with previous studies that observed wild caught hybrids [36,68]. This provides direct observation of admixture in the wild.

Finally, lower levels of genome-wide divergence (both *F*_ST_ and *d*_XY_) between sympatric pairs compared to allopatric pairs also indicate the presence of gene flow. Our *G*_MIN_ and *f*_d_ genome scans showed a higher number of putatively introgressed regions between *G. nipponicus* and Japanese Pacific *G. aculeatus* than between *G. nipponicus* and Atlantic *G. aculeatus*, suggesting that introgression has been occurring even after the Atlantic and Pacific stickleback populations diverged approximately 400,000 years BP. Our partitioned D statistics demonstrated that gene flow from *G. nipponicus* into Japanese Pacific *G. aculeatus* may be more predominant than the opposite direction in sympatry.

### High genomic divergence at a late stage of speciation with gene flow

Compared to young species pairs, less is known about the patterns of genomic differentiation at more advanced stages of speciation with gene flow. Our study provides insight on this under-represented stage of divergence. Our ABC analyses placed the estimated divergence time of *G. aculeatus* and *G. nipponicus* at 0.68 million years BP. Similarly, our Bayesian coalescent analysis of mitogenome divergence revealed a 1.3 million year split between the Japanese and Atlantic-Pacific *Gasterosteus* mitochondrial clades. An older divergence time is somewhat expected from the mitochondrial genome, given its fourfold lower effective population size [48]. Because splitting of mitochondrial lineages can occur by geographical structuring without speciation [69,70], our estimate of mitochondrial divergence may reflect more ancient geographical structuring that occurred prior to species divergence. Nonetheless, both mitochondrial and nuclear split estimates suggest that divergence between *G. aculeatus* and *G. nipponicus* occurred well before the end of the last glacial period. Therefore the Japanese stickleback system is older than all other previously examined postglacial sympatric or parapatric species pairs, which have typically diverged within the last 20,000 years [30].

The Japanese stickleback system also has a mean genome-wide *F_st_* value of 0.628, higher than any other sympatric or parapatric stickleback species pair studied so far (Fig 3C); placing this pair at the furthest end of the speciation continuum. A recent meta-analysis quantifying genetic differentiation pointed to a lack of evidence of sympatric or parapatric species pairs with *F*_ST_ between 0.3 and 0.7[8]. The Japanese species pair is therefore an example of an underepresented sympatric species pair that maintains high divergence in the face of gene flow. Within the stickleback species complex, this high divergence is in stark contrast to the typical low baseline divergence interspersed with regions of high differentiation observed in more commonly studied species pairs such as lake-stream or freshwater-anadromous pairs [16,33,71].

The primary explanation for the observed elevated divergence is most likely the more ancient divergence time of the Japan Sea-Pacific Ocean species pair compared to postglacial species pairs [16,72], However, the results of our demographic analyses indicate that high divergence is not due to a long period of allopatric isolation without gene flow, contrary to what has previously been suggested [37,39], This is important, as failing to account for variation in evolutionary history among species pairs placed on a continuum will obscure the processes leading to higher differentiation as speciation progresses. A further explanation for the high genomic divergence is the presence of strong isolating barriers between the Japan Sea and Pacific Ocean sticklebacks. Total reproductive isolation (0.970) is greater than in all postglacial species pairs (0.716-0.895) [43] and arises from a combination of habitat [41,42], temporal [62] and sexual isolation, and hybrid sterility [36,37], Recent theoretical studies have shown that selection on many barrier loci in the face of gene flow may result in a transition from low to high differentiation as a result of‘genome-wide congealing’ [10,19], It is important to note that we lack evidence that such a transition might explain the high differentiation we see here relative to the rest of the stickleback continuum (Fig 3C). However, pervasive selection on multiple isolating barriers in the Japanese stickleback system does suggest that this process could have contributed to the high genomic differentiation we observe [5].

### Localized introgression at a late stage of speciation with gene flow

Our study has also demonstrated two important signatures of introgression in the Japanese sympatric stickleback pair. Firstly, levels of background genome differentiation between *G. aculeatus* and *G. nipponicus* estimated by *F*_ST_ were lower in sympatry compared to allopatry. The higher overall genetic differentiation between *G. nipponicus* and Atlantic *G. aculeatus* is likely due to genetic drift and local adaptation and the fact that these two lineages have never overlapped geographically. Secondly and strikingly, we identified small regions of localised introgression dispersed throughout the genome when *G. nipponicus* and *G. aculeatus* co-occur in sympatry. These introgression regions were measured using *G*_MIN_, the ratio of minimum *d*_XY_ to mean *d*_XY_, where low values indicate introgression [26] and *f*_d_, the proportion of introgressed sites in a genome window [54].

Several methodological issues might influence these measures of introgression. Firstly, both *G*_MIN_ and *f*_d_ are sensitive to sample size; fewer individuals will mean rare haplotypes have a lower sampling probability. However by re-conducting our analyses using only females, a much smaller sample size than our main analysis, we still identified clear signals of introgression. Secondly, *G*_MIN_ will be biased downwards if a recently backcrossed individual is included in the dataset All Japanese *G. aculeatus* and *G. nipponicus* used in the study were identified as ‘pure’ individuals with genotyping at multiple microsatellite loci prior to resequencing [36,40]. Furthermore, we examined the identity of the two haplotypes producing the lowest value of *d*_XY_ in *G*_MIN_ valleys and confirmed that these came from different individuals in each case (data not shown). Nonetheless, even a single haplotype segregating in a population due to ILS can lower the value of *G*_MIN_; i.e. ILS could produce small, interspersed *G*_MIN_ valleys. However, ILS cannot explain why more introgressed regions occur in sympatry (Fig 5A) or why there is a clear association with an increased proportion of introgressed sites (i.e. *f*_d_) and *G*_MIN_ (Fig S9 & S13).

What then underlies the localised pattern of introgression we observe? One possible explanation is the fact that many isolating barriers are involved in reproductive isolation [36,43]. Although the genomic basis of these isolating barriers remains unknown, it is likely that barrier loci occur throughout the genome; pervasive selection at multiple loci is expected to limit the extent of introgression at this scale [73]. The strength and extent of selection at a barrier locus and genomic regions linked to it is proportional to recombination rate [73]. Recombination determines effective migration rate [74]; when recombination is high, neutral and adaptive loci linked to the target of negative selection in the recipient population have a greater probability of escaping removal and so their probability of introgression is greater [3]. Selection has a higher efficiency in these high recombination rate regions due to increased effective population size - therefore deleterious introgression is also more likely to be removed. The expectation then is that signatures of introgressed neutral or adaptive alleles are most likely to persist in regions of the genome where recombination rate is sufficiently high enough, and indeed, the positive association between introgressed regions and recombination rate we observed supports this (Fig 5B, Fig S15). Introgression is typically lower on sex chromosomes relative to autosomes in multiple taxa due to the effects of reduced recombination and greater exposure to selection in the hemizygous sex [75]. The sex chromosomes play an important role in the Japanese stickleback system, harbouring QTL for hybrid sterility and behavioural isolation [36]. Consistent with this, we observed lower introgression on the neo-sex chromosome (Fig 5E & F), although we cannot exclude the possibility that the fusion occurred more recently than the speciation event, so the opportunity for introgression on the neo-sex chromosomes was simply low relative to the rest of the genome. Taken together, our findings suggest that strong divergent selection and recombination rate variation may determine the localised signature of introgression in the genome.

The nature of gene flow in the Japanese stickleback system may also give some clues as to why we observe such highly localised introgression. One possibility is that a proportion of the introgression we detected is adaptive; i.e. it is maintained because of either directional or balancing selection. Adaptive introgression has been detected in a wide range of taxa [76], including humans [77], However, the expected signatures of the process remain unclear - especially when introgression is widespread in the genome, as is possibly the case here. Our GO analyses suggest an enrichment of immune response genes, including MHC genes, and metabolism genes in introgressed regions. Immune genes have been identified as being under balancing selection between hybridising taxa, particularly plants [78] and birds [79]. Several genes involved in metabolism are also reported to be under balancing selection in humans [80]. Furthermore, recent analysis suggests that negative frequency dependent selection might result in introgression of rare MHC alleles between divergent stickleback ecotypes [81]. Further research is necessary to directly test whether this process might explain introgression in the Japanese stickleback system.

## Conclusion

Much of our knowledge of how genomic differentiation builds along the speciation continuum is drawn from studies focusing on young, allopatric or completely reproductively isolated species pairs. Very few examples of species pairs at a later stage of divergence with on-going gene flow have been investigated. Here, we have shown that the Japan Sea and Pacific Ocean species pair exemplifies this under-represented stage of speciation and is situated at the further end of the stickleback species continuum. The high genomic differentiation between the species may be due to a more ancient divergence time than previously studied postglacial species pairs, selection on multiple isolating barriers or a combination of the two. Despite high differentiation, gene flow is on-going between the species and we identified localized signatures of introgression throughout the genome. Although the localized nature of the introgression remains unclear, selection - either directional or balancing - may play some role in promoting it Overall, our study demonstrates that high levels of genomic divergence can be established and maintained in the presence of gene flow. Further genomic studies on more species pairs at late stages of speciation with gene flow will help to understand the generality of the patterns seen here.

## Materials and Methods

### Ethics Statement

All animal experiments were approved by the institutional animal care and use committee of the National Institute of Genetics (23-15, 24-15, 25-18).

### Sample collection, whole genome resequencing and RAD sequencing

Collection and sequencing of all Japanese individuals used for whole genome resequencing has been described previously [40] except the allopatric Japan Sea fish. Briefly sympatric populations were captured from the Akkeshi system in Hokkaido, Japan in 2006 (Fig 1C). The allopatric Japan Sea female was collected in Lake Shinji in March 2014. The outgroup species, *G. wheatlandi* was captured from Demarest Lloyd State Park, MA, USA in 2007, as described previously [40]. Libraries were constructed with TruSeq DNA Sample Preparation Kit (Illumina) and whole-genome 100 bp paired-end sequencing was performed on an Illumina HiSeq2000 at the National Institute of Genetics (sympatric JS and PO) and Functional Genomics Facility, NIBB Core Research Facilities (allopatric JS) [40]. Whole genome sequencing of North American marine and stream populations collected from Little Campbell River, BC, Canada was reported previously [52,82], For the six Atlantic *G. aculeatus* individuals (North Sea) included in the study, we used previously published sequences [46].

Japanese individuals used for RAD sequencing have been previously described elsewhere [58], Samples used for RAD sequencing from the Atlantic lineage were collected from across Ireland in 2009-2011 [70,83], DNA was extracted using a Qiagen DNeasy Blood and Tissue Kit (Qiagen, Valencia, CA, USA). Single digest RAD-sequencing was performed using *Sbfl* following a standard protocol [84], RAD library preparation and sequencing was conducted using a lOObp single-end Illumina HiSeq by Floragenex (Oregon, USA).

Accession numbers, sample names and locations for all genome and RAD-seq samples are listed in Table S6.

### Whole genome alignment and variant calling

Sequence reads were mapped to the stickleback reference genome using CLC Genome 8.0 as described previously [40]. Alignments were exported as bam files and were sorted and indexed using samtools 1.2 [85]. Variant calling was carried out in two different phases. The first phase was used to call consensus bases at all sites (i.e. variant and invariant) across the genome for all 27 resequenced individuals and the outgroup (G. *wheatlandi*). Mapped reads from all individuals were piled-up using samtools *mpileup* and called against the stickleback BROAD reference genome using the bcftools 1.2 consensus caller without any filters. This consensus call produced a vcf file with a base call for every position in the genome for all samples (27 + 1 outgroup). Consensus calls from this phase were used in later demographic inference using PSMC and ABC. The aim of the second, more stringent variant calling phase was to produce a subset of high-quality polymorphic SNPs with which to examine genome-wide differentiation between the Japan Sea, Pacific and Atlantic Ocean lineages. We used bcftools to filter the consensus-call vcf for these three lineages, only retaining sites with a Phred Quality score >10, and with a maximum individual read depth of 200. Additional downstream filters were applied prior to estimating population genomic parameters from this subset vcf (see below).

### Mitochondrial genome divergence

To estimate divergence times based on mitochondrial DNA, we performed Bayesian coalescent analysis using BEAST v2.2.1 [86]. From our resequencing data, we extracted the whole mitochondrial genome from the 26 Japan Sea, Pacific and Atlantic Ocean individuals. We also downloaded two *G. wheatlandi* whole mitogenomes as outgroups (NCBI accession numbers: AB445129 & NC011570). Note that due to poor sequence coverage across the mitogenome we excluded our own re-sequenced *G. wheatlandi* individual here. Mitogenomes were aligned using MUSCLE v3.81.3 [87] resulting in a 16,549 bp final alignment.

Although there is a considerable three-spined stickleback fossil record, it is unfortunately of little use for providing fossil calibration dates for splits within the *Gasterosteus* genus [88,89]. However biogeographical events can also be used to calibrate node estimates and as such we used a normal prior (mean = 1.5 million years, SD = 0.75 million years) on the split between the Japan Sea and Pacific Ocean *G.aculeatus* lineages. We provided a further normal prior on the split date between the Pacific and Atlantic Ocean mitochondrial lineages (mean = 0. 5 million years, SD = 0.25 million years). The latter prior distribution was intentionally made wide to reflect uncertainty surrounding this estimate. Initial analyses with BEAST indicated that marginal prior distributions for node ages did not behave as specified in the model and instead returned extremely recent divergence times with low likelihood support. This is a common bias in coalescent divergence time dating and use of a calibrated prior removed this issue [90,91]. As a result, we performed all further analyses with a calibrated Yule prior. Incorrect choice of molecular clock model can seriously bias coalescent estimates of lineage divergence times and so care must be taken to ensure the appropriate model is chosen [92,93]. We used path-sampling analysis in BEAST to estimate model marginal likelihoods for three different clock models - strict, relaxed lognormal and relaxed exponential. For each model, Markov chain Monte Carlo (MCMC) was run for 5 × 10^7^ with 60 steps and marginal likelihoods were calculated using BEAST. We then ran the final model using two 10^8^ independent MCMC runs. Runs were assessed in TRACER [94] to ensure convergence and that ESS values > 200 - i.e. the posterior was adequately sampled. Independent runs were then combined to produce posterior estimates of divergence times and substitution rates.

### Nuclear phylogenetic analysis and genealogical sorting index (*gsi*)

To investigate nuclear phylogenetic discordance, we constructed maximum likelihood trees from consensus sequences for non-overlapping 10, 50 and 100 kb sliding windows following Martin et al [28]. The best-fit tree was estimated for each window using RAxML with a ‘GTRGAMMA’ model and a random number seed [95]. Trees were classified using a custom R script https://github.com/markravinet) that binned trees based on whether they matched three different topologies; species, geography, ancestral - or were unresolved. For the species category, all Atlantic, Pacific and Japan Sea individuals form separate monophyletic groups; for the geography category, Japan Sea and Pacific Ocean form a monophyletic group separate to the Atlantic Ocean; trees where the Atlantic Ocean grouped monophyletically with the Japan Sea were classed as ancestral. Trees that did not fit any of these categories were classified as unresolved. Following categorisation, trees were then standardised to ensure equal branch lengths using the *compute.brlen* function from the *Phytools* R package [96] and were finally visualised for each gene tree class using the *densiTree* function in the R package *Phangorn* [97].

We additionally used the non-overlapping Maximum Likelihood phylogenies to calculate genealogical sorting index *(gsi*) [47], We used a custom R script to estimate (*gsi)* across the autosome of 26 resequenced individuals. This allowed us to compare autosomal signals of introgression with a reduction in *gsi*.

### Population size change over time

We used PSMC to estimate fluctuations in effective population size over time [98]. PSMC uses the density of heterozygote sites across a single diploid genome to estimate blocks of constant TMRCA that are split by recombination and then uses these to infer ancestral effective population sizes (*N*_e_) over time [61,98]. Since PSMC can only analyse a single genome at a time, we ran the program separately on each of the 26 resequenced genomes from Japan Sea, Pacific and Atlantic Ocean lineages. We additionally ran the analyses for a reseqeuenced genome of a marine ecotype fish from Little Campbell River, Canada as representatives of the Eastern Pacific. Consensus sequences for each genome were converted to PSMC format - a binary format indicating the presence/absence of heterozygous sites within a specified window. We used 100 bp windows, requiring a minimum of 10,000 ‘good’ sites to be present on a genome scaffold in order for it to be included; heterozygous sites with a Phred Quality score <20 were removed. We then ran PSMC for 30 iterations with a maximum coalescent time of 15 (measured in units of *2No* where *No* is ancestral population size). Due to the difficulty of inferring past effective population sizes across this time, PSMC requires the user to provide intervals which are combined to produce the same effective population size [98]. Since this method is least accurate for recent (i.e. < 20 kyr BP) and more ancient periods [98], we estimated *N*_*e*_ for 47 intervals, combining the first four and the last three using the command “4+19*2+4”. To scale our results from coalescent units, we assumed a generation time of 1 year [99] and used an autosomal substitution rate of 7.1 × 10^-9^ [100]. Finally, to provide confidence intervals for our *N*_*e*_ estimates, we performed 100 bootstraps on 500 kb segments for each analysis.

### Approximate Bayesian Computation (ABC)

We used ABC to test different scenarios of divergence between the Japan Sea and Pacific Ocean lineage and to estimate demographic parameters, such as divergence time and migration rate, under these scenarios.

To obtain loci suitable for our ABC analysis, we randomly sampled nuclear loci from the 20 resequenced genomes (sympatric Japan Sea and Pacific Ocean) using a similar approach to Nadachowska-Brzyska et al [14]. Using a custom R script, we produced a bed file of reference genome coordinates for 2 kb loci randomly sampled at 125 kb intervals; resulting in 2,378 loci potential per individual. Using a custom python script, we called consensus sequences for each locus from the consensus vcf. This script created two haplotype sequences for each of the 2 kb loci, randomly assigning heterozygous variants to one of the two called haplotypes; this step allowed us to use unphased data for demographic analyses [66,101]. We then further filtered these loci to include only those that occur on autosomes, with >1,000 bp sequence and a minimum of 30% coverage (i.e. > 70% bases were called) for all 20 individuals. This resulted in a final dataset of 1,874 loci. Functions and scripts for generating coordinates and extracting and filtering consensus sequences are available on GitHub https://github.com/markravinet/genomesampler)

Following Robinson etal [66] we used a custom R-based control script and msABC [102] to perform simulations, calculate summary statistics and quantify their distribution across the genome in a single step. This approach offers considerable flexibility in establishing prior probability distributions for each of the estimated parameters. Furthermore, given the large size of our dataset (i.e. approximately 2,000 loci for 20 individuals) each simulation produces a large amount of data, making storage a challenge. Using R to interface with msABC allowed us to greatly reduce the required data storage.

For each of the 15 models we performed 10^6^ simulations. We used a combination of GNU Parallel [103] and independent runs across multiple computing cores to reduce analysis speed to approximately 1 day per model (scripts and additional instructions available on Github: https://github.com/markravinet).

We initially ran our simulations to produce all the available summary statistics that msABC calculates. However since summary statistic choice can greatly alter the outcomes of ABC analyses [104,105], all post-simulation ABC analyses were conducted using subsets of 29, 20 and 12 summary statistics. Following completion of the simulation step, we performed a neural-network rejection step on log-transformed parameter estimates with a tolerance of 0.01 using the abc function in the R package *abc* [106]. Posterior probability was estimated for each model using the R *abc postpr* function with multinomial logistic regression for a range of tolerance values representing 0.1%, 0.5%, 1% and 3% of the simulated data (i.e. 1,000, 5,000,10,000 and 30,000 datasets respectively). In keeping with a hierarchical analysis [14], we performed two rounds of model selection. We first chose the growth model with the highest posterior probability within each divergence scenario. Following this, we performed model selection on the five models with the highest support within each divergence category.

In order to ensure our ABC approach was reliable, we used pseudo-observed datasets (PODs) to assess how well we could discriminate between different divergence scenarios. Essentially, this involves randomly selecting a series of simulated dataset from a known model (hence pseudo-observed) and then rerunning the model selection procedure to see whether the true model could be recovered. For further details of our POD-based sensitivity analysis and ABC approach, see the *Supplementary methods* section.

### Detecting genome-wide divergence and recent introgression

Weir and Cockerham’s *F*_ST_ [107] was calculated using 10 and 50 kb non-overlapping windows with VCFtools 0.113 [108]. To calculate haplotype-based statistics such as *d*_XY_, *G*_MIN_ and *f*_d_, we used a modified version of a python script used by Martin et al [28]. In addition to our main filters on the dataset (see *Genome Alignment and Variant calling*), we only calculated these haplotype-based statistics for windows with >50% of useable bases - i.e. >5,000 sites within a 10 kb sliding window. For autosomal statistics, all individuals were included in the analyses. For comparing the ancestral (chrXIX) and neo-sex chromosomes (chrIX) with autosomes (Fig. 5E and 5F), we re-ran the analyses of all chromosomes using only females. In addition to 10 kb windows, we also performed analyses for non-overlapping 50 kb windows to aid visualisation; the results from all analyses were then combined into a single dataset using custom R scripts.

We calculated recently established statistics, *G*_MIN_ and *f*_d_, for detecting introgression between divergent lineages [26,54], G_MIN_ IS particularly suited for identifying recent, low frequency introgression [26] whereas *f*_d_ can also identify stronger, high frequency introgression events [54], Importantly, both methods are robust to variation in recombination rate variation. Initial genome scans conducted using G_*MIN*_ revealed a series of‘valleys’ present across the genome. Detection of such valleys, like genomic islands of divergence, presents a variety of methodological issues. Firstly, how do we determine that G_*MIN*_ valleys are not due to stochastic variation in genealogy amongst loci? Secondly, how do we measure the size and distribution of valleys of introgression? Finally, how can we determine a null or expected distribution of valleys across the genome to test for the under- or overrepresentation of valleys?

To deal with each of these issues in turn, we first performed chromosome-specific permutations to identify the null distribution of the value of G_*MIN*_. Specifically, we shuffled the nucleotide sequence of each chromosome 100 times and estimated G_*MIN*_ for 11 different sliding window sizes (5,000, 5,500, 6,000, 6,500, 7,000, 7,500, 8,000, 8,500, 9,000, 9,500 and 10,000 kb), representing the distribution of useable sites from the empirical dataset We then used the lower 99 percentile of the permutations to determine the mean value of G_*MIN*_ below which a window could be classified as a valley. Identifying the boundaries of divergent genome regions is somewhat subjective and open to potential bias [109]. To account for this, we used a hidden Markov-model (HMM) approach to classify windows into two states - i.e. valleys or non-valleys - and to estimate the probability of state transition. Following Soria-Caracasco *et al*. [55], we used the R package HiddenMarkov [110] on a logit transformed G_*MIN*_ distribution. Transition probabilities between the two states were symmetrical with an emphasis on it being difficult to transition between states (p = 0.1) but relatively easy to remain within a state (p = 0.9). Since valleys are relatively rare in the genome, we set our models to start in the non-valley state and we provided estimated parameter values for the states based on the empirical distribution. Our permutation test to find the 99 percentile therefore also provided an empirical basis for identifying valleys. HMM estimates were run for both the sympatric and allopatric comparisons using the *baumwelch* function to estimate parameters using the Baum-Welch algorithm and the *viterbi* function to estimate the sequence of states using the Viterbi algorithm. We used a similar approach to identify *f*_d_ peaks but we instead performed the analysis using untransformed *f*_d_ values only in the sympatric Japanese *G. aculeatus* and *G. nipponicus* comparison.

### RAD-seq data processing and admixture analysis

RAD sequence reads were demultiplexed and processed using the *process_radtags* module of Stacks 1.30 [111]. All reads were trimmed to 90 bp and any read where the average Phred quality score dropped below 10 in a 9 bp sliding window was discarded. Following filtering, reads were mapped to the Roesti *et al*. [59] build of the *G. aculeatus* genome using GSNAP [112] allowing a maximum of two indels to be present in an alignment, reporting no suboptimal hits, allowing a maximum of 8 mismatches and printing only the best alignment. SNPs were then called using the samtools and bcftools pipeline [113]. Called variants were then filtered using vcftools to remove all sites with greater than 25% missing data, to include genotypes only with an individual depth between 15X and 100X, to remove all sites with a Phred quality score below 20 and with a minor allele frequency below 0.05. Since common admixture analyses assume independence among sites (i.e. the absence of linkage disequilibrium) [114], we additionally pruned our RAD-derived SNP dataset using plink [115], removing all sites where pairwise linkage disequilibrium was greater than 0.4.

PCA on allele frequencies from all individuals was conducted using the *glPca* function from the R package adegenet [116]. Admixture analysis was carried out on a subset of samples from the Japanese archipelago using STRUCTURE [50,51]. For each value of *K* from 1 to 8, the program was run for 10 iterations with a burn-in of 10,000 steps followed by 20,000 MCMC steps. The most likely value of *K* was assessed using STRUCTURE HARVESTER [117].

### Detecting the direction of introgression

We investigated the direction of gene flow between the Japan Sea and Pacific Ocean lineages using partitioned *D* statistics [56]. This is conceptually similar to standard four population ABBA-BABA tests for gene flow but includes a fifth population - an allopatric lineage of the Japan Sea. This balances the assumed phylogeny (i.e. ((PI, P2), (P3_1_, P3_2_), 0) and therefore allows us to rotate the populations used in the analysis - i.e. testing for an enrichment of gene flow in both directions. We therefore tested two topologies ((AT, PO), (JS_s_, JS_A_), 0) and ((JS_A_, JS_s_), (PO, AT), 0) (see Fig S14). For either test topology, an excess of the ABBAA (compared to BABAA) or ABBBA (compared to BABBA) positions in a genome window inflates partitioned D statistics above zero - indicating gene flow from the P3 into P2.

Given that the partitioned *D* approach has attracted some criticism, we also calculated *D*_FOIL_ statistics [57], *D*_FOIL_ is an additional extension of the four population test but one that incorporates all possible introgression events for a symmetric four population tree (excluding the outgroup). We used the same test phylogeny as with the partitioned *D* statistics (see Figs S14 &S15).

Both partitioned *D* and *D*_FOIL_ based on ABBA/BABA methods - i.e. where only a single individual is present at the tips of the phylogeny. To account for this, we extended both methods to account for allele frequency data, meaning our site pattern counts are weighted by allele frequencies [54], To calculate both *D* and *D*_FOIL_ statistics, we used a modified version of a python script used by Martin etal [28].

### Characterization of introgression sites

In order to characterize regions of introgression, we identified candidate regions showing a strong signature of introgression (i.e. G_*MIN*_ valleys and *f*_d_ peaks) from our genome scan approach. We then counted the number of unique genes falling within our candidate valleys/peaks and compared this to a null distribution generated by 1,000 random samples of 10 kb non-valley/non-peak genome windows for the same number and size range as the valleys or peaks.

We then tested whether genes in introgressed regions were more likely to have any specific functions. To achieve this, we used gene ontology (GO) analysis on genes in valleys and 1,000 randomly chosen from across the genome. GO analysis was performed with the ClueGO plugin [118] for Cytoscape 3.4.0 [119]. Since functional annotations for this analysis were drawn from the human genome, we first generated a list of human-stickleback orthologous gene IDs (Ensembl Biomart 86). We then subset our candidate and random gene sets to include only orthologous genes. Several human genes have multiple stickleback orthologs; we therefore allowed only a single, randomly chosen occurrence of each human gene in both sets to prevent pseudo-replication. A hypergeometric test was conducted for testing enrichment with Benjamini & Hochberg FDR correction [120].

## Acknowledgements

We are grateful to Manabu Kume, Seiichi Mori, and staff at the Aquarium Gobius for providing samples and Katsushi Yamaguchi for technical assistance. Keisuke Honda, the Institute of Statistical Mathematics and the DNA Data Bank of Japan are thanked for their help with running analyses on supercomputers. We are additionally grateful to Simon Martin, John Robinson and David Marques for sharing scripts and their advice on analyses. Freddy Chain and Philine Feulner are also thanked for their assistance with their sequence data. We thank Cassandra Trier for her assistance and advice with GO analyses. All members of the Kitano Lab provided invaluable advice throughout the project. We would also like to thank Mark Kirkpatrick and members of his lab for comments on manuscript.

## Supporting information

**S1 Table. Classification of trees.** Proportion of trees drawn from 10, 50 and 100 kb windows representing species, ancestral and geographical topologies.

**S2 Table. Mitogenome analysis.** Marginal log-likelihood values and Bayes factor comparisons from path sampling of substitution clock models used for mitogenome phylogeny and divergence time estimation.

**S3. Table. Median, L95% and U95% HPD (highest probability densities) for demographic parameters estimated under IM + bottleneck model.**

**S4 Table. Number of valleys and valley per Mb for all chromosomes.**

**S5 Table. Enriched GO terms for genes present in *G*_MIN_ valleys.**

**S6 Table. Enriched GO terms for genes present in *f*_d_ peaks.**

**S7 Table. Sample names, locations and accession numbers.**

**S1 Fig. Global species distribution.** The global distribution of the Pacific and Atlantic Ocean lineages of *G. aculeatus* allow sympatric and allopatric comparisons with *G. nipponicu;* AT = Atlantic Ocean, PO = Pacific Ocean and JS = Japan Sea.

**S2 Fig. Japanese stickleback mitonuclear discordance.** (A) Mitogenome Bayesian tree shows divergence between two mitochondrial clades - the Transpacific (TPN) and European North American (ENA); asterisks on nodes indicate appropriate densities shown in (B). (B) Posterior probability densities for mitochondrial divergence time between *G. aculeatus* and *G. nipponicus* and between Pacific and Atlantic populations of the ENA clade.

**S3 Fig. Bootstrapped PSMC curves for 26 resequenced individuals.**

**S4 Fig. PSMC profile for all 26 individuals and an additional Eastern Pacific individual from Little Campbell River, Canada.**

**S5. Fig. Principal component analysis on RAD-seq data from 295 individuals from across the distribution of all three lineages.** The arrow indicates the presence of an admixed individual occurring in the Akkeshi system.

**S6 Fig. STRUCTURE analysis on RAD-seq data from Japanese populations.** Analysis with *K=* 2 clusters (A), which is supported by likelihood analysis (B), showed the presence of admixed individuals in the Akkeshi system.

**S7 Fig. Genome-wide *F*_ST_ measured in non-overlapping 50 kb windows for allopatric and sympatric between and within species comparisons.**

**S8 Fig. Genome-wide *d*_XY_ measured in non-overlapping 50 kb windows for allopatric and sympatric between and within species comparisons.**

**S9 Fig. Negative association between *f*_*d*_ and *G*_MIN_ in sympatric (JS vs PO) and allopatric (JS v AT) between species comparisons.**

**S10 Fig. Genome-wide G_MIN_ for sympatric between species comparisons.** Black line represents 50 kb non-overlapping window *G*_MIN_ signature. Points represent 10 kb windows; grey points are non-valley windows, blue points are valley windows identified by Hidden Markov Model algorithm.

**S11 Fig. Absolute divergence [*d*_XY_] is lower and *f*_d_ is higher in G_MIN_ valleys compared to the genome-wide background.**

**S12 Fig. Genome-wide *f*_*d*_ for sympatric between species comparisons.** Black line represents 50 kb non-overlapping window *f*_d_ signature. Points represent 10 kb windows; grey points are non-valley windows, blue points are peak windows identified by Hidden Markov Model algorithm.

**S13 Fig. Comparison of *d*_XY_ between *f*_d_ peaks and non-peaks.**

**S14 Fig. Analysis of partitioned *D* statistics.** (A) Boxplots comparing partitioned *D* statistics between *f*_d_ peaks and the autosomal background. Dashed line at zero indicates a balance between allele patterns indicative of incomplete lineage sorting. In (B), P1 = Atlantic Ocean (AT), P2 = Pacific Ocean (PO), P3_1_ = sympatric Japan Sea (JSs), P3_2_ = allopatric Japan Sea (JS_A_), and 0 = *G. wheatlandi* (WT). In (C), P12 and P3 were swapped. Di measures asymmetry between PI and P2 where the derived allele B is present in P3_1_ but not P3_2_, D2 measures where allele B is present in P3_2_ but not P3_1_, and D12 measures where the derived allele is shared by both P3i and P32. If we assume that the derived allele B occurred at the ancestor of P3, D12 indicates introgression from P3 to P2. See [56] for a more detailed explanation of these statistics.

**S15 Fig *D*_FOIL_ statistics.** In this statistics, we assume that the divergence time between P_1_ and P_2_ is younger than that between P_3_ and P_4_ and infer all possible introgressions including ancestral introgression involving P_12_. The number of loci (100 kb-window) that show statistically significant introgression are shown. See [57] for a more detailed explanation of these statistics.

**S16 Fig. The relationship between introgression measured as *G*_MIN_ and *f*_d_ and log_10_ recombination rate.**

**S17 Fig. Venn diagram showing overlap between genes occurring in introgressed regions identified using different measures.**

**Supplementary methods - ABC analysis.**

